# Thermal acclimation of photosynthetic activity and Rubisco content in two hybrid poplar clones

**DOI:** 10.1101/438069

**Authors:** Lahcen Benomar, Mohamed. Taha Moutaoufik, Raed Elferjani, Nathalie Isabel, Annie DesRochers, Ahmed El Guellab, Rim Khlifa, Lala Amina Idrissi Hassania

## Abstract

The mechanistic bases of thermal acclimation of net photosynthetic rate (*A*_*n*_) are still difficult to discern and empirical research remains limited, particularly for hybrid poplar. In the present study, we examined the contribution of a number of biochemical and biophysical traits on thermal acclimation of *A*_*n*_ for two hybrid poplar clones. We grew cuttings of *Populus maximowiczii* × *Populus nigra* (M×N) and *Populus maximowiczii* × *Populus balsamifera* (M×B) clones under two day/night temperature of 23°C/18°C and 33°C /27°C and under low and high soil nitrogen level. After 10 weeks, we measured leaf RuBisCO and RuBisCO activase (*RCA*) amounts and the temperature response of *A*_*n*_, dark respiration (*R*_d_), stomatal conductance, (*g*_s_), maximum carboxylation rate of CO_2_ (*V*_cmax_) and photosynthetic electron transport rate (*J*). Results showed that a 10°C increase in growth temperature resulted in a shift in thermal optimum (*T*_*opt*_) of *A*_*n*_ of 6.2±1.6 °C and 8.0±1.2 °C for clone M×B and M×N respectively, and an increased *A*_*n*_ and *g*_s_ at the growth temperature for clone M×B but not M×N. RuBisCO amount was increased by N level but was insensitive to growth temperature while *RCA* amount and the ratio of its short to long isoform was stimulated by warm condition for clone M×N and at low N for clone M×B. The activation energy of *V*_*cmax*_ and *J* decreased under warm condition for clone M×B and remain unchanged for clone M×N. Our study demonstrated the involvement of both *RCA*, activation energy of *V*_*cmax*_ and stomatal conductance in thermal acclimation of *A*_*n*_.

## Introduction

Global warming may lead to a significant reduction of forest productivity through a decrease in net assimilation rate of CO_2_ (Lloyd and Farquhar, 2008; Sage et al., 2008). Plant physiological processes including light-saturated photosynthetic rate (*A*_*n*_) and dark respiration (*R*_d_) are strongly temperature-dependent and their acclimation may help trees maintain a normal growth when temperature shifts from optimum to warm (Atkin et al., 2005; Medlyn et al., 2002; Sage et al., 2008). Thermal acclimation of *A*_*n*_ is achieved through adjustments of one or more morphological, biochemical and biophysical components of photosynthesis which may occur via (i) a shift of the thermal optimum of *A*_n_ (*T*_*opt*_) toward the new growth temperature (ii) an increase or a maintenance of the photosynthetic rate at *T*_*opt*_ (*A*_*opt*_) at warmer growth temperatures (iii) a shift in both *A*_*opt*_ and *T*_*opt*_, and (iv) an increase or a maintenance of the photosynthetic rate respective to growth temperature (*A*_*growth*_) (Sage and Kubien, 2007; Way and Yamori, 2014; Yamori et al., 2014). The mechanisms involved in thermal acclimation of photosynthesis are still difficult to discern and may originate, among others, from species thermal origin (Yamori et al., 2009). They include modulation of (i) basal maximum carboxylation rate *V*_cmax_^25^ or maximum electron transport rate *J*_max_^25^ (measured at reference temperature of 25°C), (ii) thermal response of both *V*_cmax_ and *J*_max_ (activation and deactivation energy), (iii) nitrogen allocation to carboxylation *vs.* elctron transport (ratio of *J*_max_ to *V*_cmax_) and (iv) thermal response of stomatal and mesophyll conductance (Hikosaka et al., 2006; Sage and Kubien, 2007; Way and Yamori, 2014; Yamori et al., 2014).

Leaf nitrogen (N) plays a key role in carbon assimilation processes and hence plant growth and survival (DesRochers et al., 2003; Fisichelli et al., 2015), as most of the leaf nitrogen is allocated to proteins involved in light harvesting, Calvin-Benson cycle and electron transfer along thylakoid membranes (Field, 1983; Poorter et al., 2009). Leaf nitrogen content is generally deficient in temperate and boreal regions and has been shown to decrease in response to increasing growth temperature (Reich and Oleksyn, 2004; Gunderson et al., 2010; Scafaro et al., 2016). A decrease in leaf N in response to increasing growth temperature may result in a decrease of RuBisCO content (Scafaro et al., 2016). This has been proposed as an explanation of the commonly observed deacrease in *V*_cmax_ at temperatures above the optimum and the resulting lack of thermal acclimation of *A*_*n*_ (Scafaro et al., 2016; Crous et al., 2018). On the other hand, Yamori et al., (2011) found that photosynthesis temperature response of several C_3_ plants was generally RuBP carboxylation-limited above the *T*_*opt*_ at low leaf nitrogen content while, under high N level, it shifted to a limitation by RuBP regeneration. However, the effect of temperature on the limiting steps of *A*_*n*_ (*V*_cmax_ *vs. J*_max_) may depend on the reponse of CO_2_ conductance (*g*_*s*_ and *g*_*m*_) as well (Benomar et al., 2018; Qiu et al., 2017; von Caemmerer and Evans, 2015; Warren, 2008). Moreover, RuBisCO-related effect on *A*_*n*_ at above-optimal temperature may depend on the plasticity of *J*_max_^25^ to *V*_*cmax*_^*25*^ ratio. From this perspective, this may be applicable only for cold-adapted plant species, which are characterized by a higher *J*_max_^25^ to *V*_cmax_^25^ ratio and low or lack of its adjustment in response to both N level and growth temperature (Benomar et al., 2018; Kattge and Knorr, 2007). Weston et al. (2007) did not observe any change in RuBisCO concentration for two genotypes of *Acer rubrum* grown under hot and optimal temperatures. Then, more research is needed to unravel the multiple factors involved in the response of carbon assimilation to above-optimal temperatures. In fact, it has been proven that *V*_cmax_ do not only depend on RuBisCO concentration but also on its activation state (inhibited/activated) (Cen and Sage, 2005; Sage et al., 2008; Salvucci and Crafts-Brandner, 2004). The activation state of RuBisCO is regulated by the RuBisCO activase (*RCA*), a heat-labile enzyme using eneregy via ATP hydrolysis to release inhibitors from the active site of RuBisCO (Crafts-Brandner and Salvucci, 2000; Salvucci and Crafts-Brandner, 2004; Yamori and von Caemmerer, 2009). A decrease in *RCA* activity has been documented as a primary cause of reducing RuBisCO activity and then photosynthetic performance in response to increasing growth temperature (Hozain et al., 2009; Salvucci and Crafts-Brandner, 2004; Yamori and von Caemmerer, 2009). *RCA* is a stromal protein existing in two isoforms of 41–43 kDa (short isoform) and 45–46 kDa (long isoform) that arises from one single gene with alternatively spliced transcript or from two separate genes. Still, the specific physiological role of a given isoform with respect to heat stress is generally not understood. Recent studies from herbaceous species demonstrated an increase in the two *RCA* forms or a shift in the balance between them when plants were exposed to temperature above 30°C (Law et al., 2001; Ristic et al., 2009; Wang et al., 2010; Weston et al., 2007; Yamori et al., 2014).

Here we used *Populus* to study the physiological thermal acclimation because of its commercial and environmental importance in the northern hemisphere and its fast growth rate. Information on the response of photosynthesis to higher temperature for tree species is limited in general, and previous studies conducted on *Populus balsamifera* (Silim et al., 2010), *Populus tremuloides* (Dillaway and Kruger 2010), *Populus nigra* (Centritto et al., 2011), *Populus grandidentata* (Gunderson et al., 2010) and *Populus deltoides × nigra*. (Ow et al., 2008) found little evidence of a thermal acclimation of *A*_*n*_ to increasing temperatures. Nevertheless, little research focused on the physiological and molecular mechanisms underlying the observed thermal acclimation of trees. The objective of the present study was to examine to what extent leaf nitrogen, RuBisCO and RCA content are involved in thermal acclimation of photosynthetic activity in hybrid poplars.

## Methodology

### Plant material and growth conditions

This experiment was conducted in greenhouses and growth chambers at Université Laval, Québec, Canada, from January to May 2017. Dormant cuttings of two hybrid poplar clones: M×N (*Populus maximowiczii* × *Populus nigra*) and M×B (*Populus maximowiczii* × *Populus balsamifera*) were provided by the Québec’s Ministère des Forêts, de la Faune et des Parcs from the forest nursery of Berthier (Berthierville, Québec, Canada) during early January after chilling needs were met. Cuttings were planted in 2 L pots filled with peat/vermiculite substrate (v/v=3/1) and placed in two greenhouses where day/night temperatures were 23°C/18°C and 33°C/27°C. Plants were grown under a photosynthetically active radiation (*PAR*) ranging between 400 and 700 μmol m^−2^ s^−1^, a relative humidity of 65% and a 8/16 h dark/light photoperiod using 400 W metal halide lamps. Cuttings were irrigated daily to maintain full soil field capacity. After 4 weeks, for a better control of growth conditions (mainly temperature and relative humidity), pots were transferred to growth chambers (model PGW 36, Conviron, Winnipeg, Canada) under a split-split-plot layout; the Temperature×Clone as first split and Nitrogen level as second split. The same environment parameters as in greenhouses were used, except PAR, which was kept at a constant rate of 500 μmol m^−2^ s^−1^ during day time. In each growth chamber, half of plants (n=18) were randomly assigned to receive a low-nitrogen fertilization treatment (5 mM) while the other half received a high-nitrogen (20 mM). Nitrogen was added, every week, using (20N-20P-20K) fertilizer dissolved in distilled water. Plants (n = 72; 2 growth temperatures × 2 nitrogen levels × 2 hybrid poplar clones × 9 replicates) were allowed to acclimate to respective growth conditions for 6 weeks before measurements were taken. Pots were moved within each chamber every third day to eliminate any position-related bias.

### Gas exchange measurements

After 10 weeks of growth, leaf-level gas exchange were measured on the 4^th^ fully expanded leaf from the top of each plant using two cross-calibrated portable open-path gas-exchange systems (Li-6400, Li-Cor Inc., Lincoln NE), equipped with a leaf chamber fluorometer (Li-6400-40, Li-Cor Inc). The measurements were made on 24 plants in total (3 replicates × 2 clones × 2 temperatures × 2 N levels). Given the limited control capacity of LI-6400 system on leaf temperature in the cuvette (*T*_*leaf*_ can be set to ± 6°C of the ambient temperature), measurements were performed in a growth chamber under controlled temperature and relative humidity. Growth chamber temperature was set manually to desired *T*_*leaf*_ allowing an effective and quick easy adjustment over the 10 - 40°C range and an exposure of the whole plant to the targeted temperature.

Temperature was increased from 10°C to 40°C with 5°C increment and plants were allowed to acclimate for at least 20 min to each step. At each temperature, we measured dark respiration (*R*_*d*_) followed by *A-C*_*i*_ response curve records with 10-minutes period between *R*_*d*_ and *A-C*_*i*_ respected to allow complete opening of stomata. *A-Ci* response curves were recorded at each temperature after at least 10 min of steady state at ambient CO_2_ partial pressure *C*_*a*_=400 μmol mol^−1^ and a saturated photosynthetic active radiation *PAR*=800 μmol m^−2^ s^−1^. The saturated *PAR* was determined from measured *A-Q* curve on 3 plants from each Clone×Growth T° combination at 25°C. Thereafter, the reference CO_2_ (*C*_*a*_) was changed in the following order: 400, 350, 300, 200, 100, 50, 400, 500, 600, 800, 900, 1000, 1200, 1400, and 1600 μmol mol^−1^. Values were recorded based on the stability of photosynthesis, stomatal conductance (*g*_*s*_), CO_2_ and water vapor concentration. The vapor pressure difference (*VPD*) during measurement varied from 0.5 to 3.2 KPa from low to high temperature and was lowered as much as possible at high temperature by maintaining relative humidity (*RH*) at 70% inside the growth chamber. Similarly, *RH* was maintained at 50% to maintain *VPD* as high as 0.5 KPa at low temperature. The list of abbreviations and symbols are given in Table 1.

**Table 1:** List of abbreviations

| Symbol | Definition | Unit |
| --- | --- | --- |
| $A_c$ | RuBP-saturated CO<sub>2</sub> assimilation rate | $\mu\text{mol CO}_2 \text{ m}^{-2} \text{ s}^{-1}$ |
| $A_{\text{growth}}$ | Photosynthetic rate at growth temperature | $\mu\text{mol CO}_2 \text{ m}^{-2} \text{ s}^{-1}$ |
| $A_n$ | Net CO<sub>2</sub> assimilation rate | $\mu\text{mol CO}_2 \text{ m}^{-2} \text{ s}^{-1}$ |
| $A_j$ | RuBP-limited CO<sub>2</sub> assimilation rate | $\mu\text{mol CO}_2 \text{ m}^{-2} \text{ s}^{-1}$ |
| $A_{\text{opt}}$ | Photosynthetic rate at $T_{\text{opt}}$ | $\mu\text{mol CO}_2 \text{ m}^{-2} \text{ s}^{-1}$ |
| $C_a$ | Atmospheric CO<sub>2</sub> concentration | $\mu\text{mol mol}^{-1}$ |
| $C_i$ | intercellular CO<sub>2</sub> concentration | $\mu\text{mol mol}^{-1}$ |
| $E_a$ | Energy of deactivation | $\text{KJ mol}^{-1}$ |
| $E_d$ | Activation energy | $\text{KJ mol}^{-1}$ |
| $g_s$ | Stomatal conductance | $\text{mol H}_2\text{O m}^{-2} \text{ s}^{-1}$ |
| $J$ | Electron transport rate | $\mu\text{mol m}^{-2} \text{ s}^{-1}$ |
| $J_{\text{max}}^{25}$ | Maximal electron transport rate at leaf temperature of 25°C | $\mu\text{mol m}^{-2} \text{ s}^{-1}$ |
| $J_{\text{max}}^{25} : V_{\text{cmax}}^{25}$ | Ratio of maximal electron transport to maximal carboxylation rate at leaf temperature of 25°C | |
| $N_{\text{area}}$ | Leaf nitrogen in area basis | $\text{g m}^{-2}$ |
| $O$ | Partial atmospheric pressure of O<sub>2</sub> | $\text{mmol mol}^{-1}$ |
| $PAR$ | Photosynthetically active radiation | $\mu\text{mol m}^{-2} \text{ s}^{-1}$ |
| $SLA$ | Specific leaf area | $\text{cm}^2 \text{ g}^{-1}$ |
| $R_{\text{day}}$ | Mitochondrial respiration in the light | $\mu\text{mol CO}_2 \text{ m}^{-2} \text{ s}^{-1}$ |
| $R_d$ | Dark respiration | $\mu\text{mol CO}_2 \text{ m}^{-2} \text{ s}^{-1}$ |
| $R_d^{10}$ | $R_d$ at leaf temperature of 10°C | $\mu\text{mol CO}_2 \text{ m}^{-2} \text{ s}^{-1}$ |
| $RCA$ | RuBisCO activase | |
| $T_{\text{opt}}$ | Thermal optimum | °C |
| $K_c$ | Michaelis–Menten constants of Rubisco for CO<sub>2</sub> | $\mu\text{mol mol}^{-1}$ |
| $K_o$ | Michaelis–Menten constants of Rubisco for O<sub>2</sub> | $\text{mmol mol}^{-1}$ |
| $Q_{10}$ | Rate of change in $R_d$ with a 10°C increase in temperature | |
| $\Gamma^*$ | CO<sub>2</sub> compensation point in the absence of mitochondrial respiration | $\mu\text{mol mol}^{-1}$ |
| $\alpha$ | Efficiency of light energy conversion | |
| $V_{\text{cmax}}$ | Maximal carboxylation rate | $\mu\text{mol CO}_2 \text{ m}^{-2} \text{ s}^{-1}$ |
| $V_{\text{cmax}}^{25}$ | Maximal carboxylation rate at leaf temperature of 25°C | $\mu\text{mol CO}_2 \text{ m}^{-2} \text{ s}^{-1}$ |

### Estimation of gas exchange parameters

The photosynthetic capacity variables, *V*_cmax_ and *J*_max_, were estimated from gas-exchange by fitting the *A-C*_*i*_ curve with the biochemical model of C_3_ (Farquhar et al., 1980), assuming infinite mesophyll conductance (*g*_m_). In fact, the estimation of *g*_m_ from *A-*Ci is very challenging as it depends on the number of data points on the *A-*Ci curve and goodness - of-fit of the curve which is difficult to achieve at high and low temperatures. In this experiment, we tried to estimate *g*_m_ from *A*-Ci curves following Ethier et al. (2004) and Miao et al. (2008) without success as about 45 % of them gave non-meaningful estimates. The model was thus fitted using non-linear regression techniques (Proc NLIN, SAS) following Dubois et al. (2007). Briefly, the net assimilation rate (*A*_*n*_) is given as:

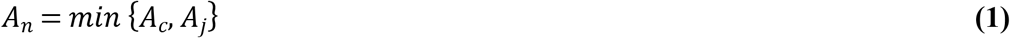

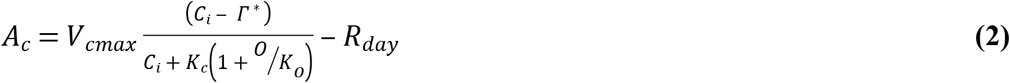

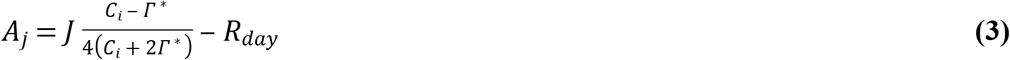

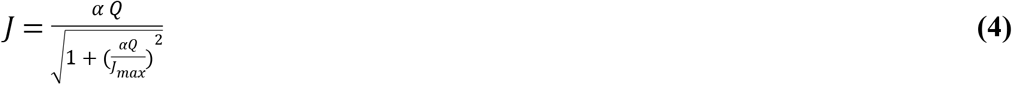

where *V*_cmax_ is the apparent maximum rate of carboxylation (*μ*mol CO_2_ m^−2^ s^−1^), *O* is the partial atmospheric pressure of O_2_ (mmol mol^−1^), *Γ\** is the CO_2_ photo-compensation point in the absence of mitochondrial respiration, *R*_*day*_, is mitochondrial respiration in the light (*μ*mol CO_2_ m^−2^ s^−1^), *C*_*i*_ is the intercellular (substomatal) concentration of CO_*2*_ (μmol mol^−1^), *K*_*c*_ (μmol mol^−1^) and *K*_*o*_ (mmol mol^−1^) are the Michaelis–Menten constants of Rubisco for CO_2_ and O_2_, respectively, *J* is the apparent rate of electron transport (*μ*mol CO_2_ m^−2^ s^−1^), *J*_max_ is the apparent maximum rate of electron transport (*μ*mol CO_2_ m^−2^ s^−1^), *Q* is the incident *PAR* (*μ*mol m^−2^ s^−1^), *α* is the efficiency of light energy conversion (0.18) which represents the initial slope of the photosynthetic light response curve (Miao et al., 2008). The values at 25°C used for *K*_*c*_, *K*_*o*_ and *Γ\** were 272 μmol mol^−1^, 166 mmol mol^−1^ and 37.4 μmol mol^−1^, respectively (Sharkey et al., 2007) and their temperature dependency were as in Sharkey et al.(2007. Most of *A-C*_*i*_ curves at 35°C and 40 °C measured for low nitrogen level at 23 °C failed to converge and estimates of *V*_*cmax*_ and *J* could not be obtained.

### Characterization of the temperature responses of gas exchange parameters

Photosynthesis temperature response curves were fitted individually with a quadratic model following Battaglia et al. (1996):

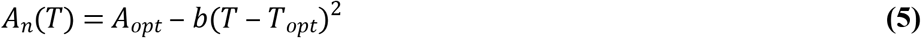

where *A*_*n*_*(T)* is the photosynthetic rate at temperature T in °C, *A*_*opt*_ is the photosynthetic rate at the temperature optimum (*T*_*opt*_) and the parameter b describes the spread of the parabola. *A*_*growth*_ was then estimated using the obtained parameters from equation (5) for each individual curve. Daytime temperature was used as growth temperature given the uncertainty regarding the effect of nighttime temperature on *A*_*n*_.

Dark respiration temperature response curves were fitted with a model in equation (6) to estimate the *Q*_*10*_ (the change in respiration with a 10°C increase in temperature) following Atkin et al. (2005):

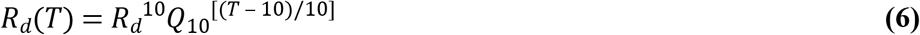

where *R*_*d*_^*10*^ is the measured basal rate of *R*_*d*_ at the reference temperature of 10°C. The responses of *V*_*cmax*_ and *J* to leaf temperature were fitted using the following two models (equation (7) and (8)) depending on the presence or not of deactivation above thermal optimum following Medlyn et al. (2002):

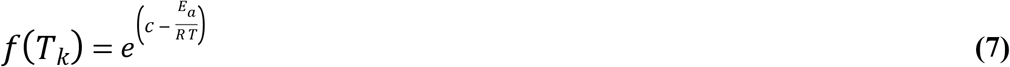

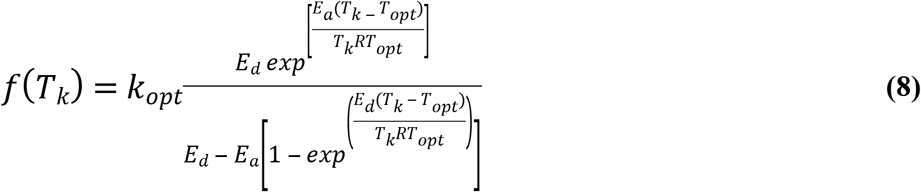

where *E*_*a*_ is the activation energy, *E*_*d*_ is the energy of deactivation, *K*_*opt*_ is the *V*_*cmax*_ or *J* at the temperature optimum (*T*_*opt*_). *E*_*d*_ was fixed at 200 KJ mol^−1^ (Medlyn et al., 2002) to reduce the number of estimated parameters to three.

### SLA and leaf nitrogen

Leaves used for gas exchange measurements were collected and immediately placed in dry ice before being stored at −20°C and processed within a week for protein extraction. The extracts were conserved under −80°C and dosage of proteins (RuBisCO and RCA) was done once all samples were extracted. Symmetric leaves (by the stem) were also collected to measure projected area with WinSeedle (Version 2007 Pro, Regent Instruments, Québec, Canada). Samples were then oven-dried for 72h at 56 °C, and their dry mass determined. Specific leaf area (*SLA*) was calculated as the ratio of the projected leaf area (cm^2^) to the leaf dry mass (g). Later, leaves were ground separately and N content determined at Université Laval using a LECO elemental analyser (LECO Corporation, St Joseph, MI, USA).

### Extraction and dosage of RuBisCO and RuBisCO activase

Proteins were extracted from frozen leaves at −20 °C within less than one week after leaf harvesting following the method outlined in Yamori and von Caemmerer (2009). Briefly, 100 mg of leaves were initially ground in liquid nitrogen using a mortar and pestle. Proteins were extracted on ice using a protein extraction buffer containing 50 mM Hepes-KOH pH 7.8, 10 mM MgCl_2_, 1 mM EDTA, 5 mM DTT, 0.1% triton X100 (v/v) and protease inhibitor cocktail (Roche). The extracts were conserved under −80°C. Once all samples were extracted, the solutions were centrifuged at 16,000g for 1 min followed by determination of the concentration of total soluble proteins (TSP) in supernatant by the Bradford method (Bradford, 1976).

After dosage, 4× sample buffer (250 mM Tris–HCl, pH 6.8, 40% glycerol, 8% SDS, 0.2% Bromophenol-blue, 200 mM DTT) was added to proteins extracts, heated at 100 °C for 5 min and then centrifuged at 16,000 g for 5 min. After cooling to room temperature, a volume representing 20 μg of total TSP extract of each sample was loaded onto 12% SDS-polyacrylamide gel electrophoresis (SDS-PAGE). The electrophoresis was carried out at room temperature at a constant voltage (120 V). Following SDS-Page, the proteins were transferred to a nitrocellulose membrane (Life Sciences, Mississauga, Canada) for western blot.

Blots were incubated with 5% non-fat milk in TBST (50 mM Tris, pH 7.5, 150 mM NaCl, 0.1% Tween-20) for 60 min, the membranes were washed twice with TBST and incubated with antibodies against RuBisCO (Agrisera AB, Vännäs, Sweden) or against RuBisCO activase (Agrisera AB, Vännäs, Sweden) at room temperature for 60 min. Membranes were washed three times with TBST for 10 min and incubated with secondary antibodies peroxidase-conjugated (Goat Anti-Chicken (abcam) for RuBisCO and Goat Anti-Rabbit (abcam) for RuBisCO activase) during 60 min at room temperature. Blots were washed with TBST three times and developed with the ECL system using Odyssey^®^ Infrared Imaging System (Li-COR, Biosciences). Images were analysed using ImageJ (Rasband, 2016) to determine band densities of each sample. The RuBisCo, *RCA* and its two isoforms concentration were expressed as relative to the sample representing the highest density (Perdomo et al., 2017; Prins et al., 2008; Ristic et al., 2009).

### Statistical analysis

Three-way analysis of variance was performed to test the effect of growth temperature, clone and nitrogen level on response variables using MIXED procedure of SAS (SAS Institute, software version 9.4, Cary, NC, USA). We used proc Glimmix for response variables (apparent *V*_cmax_^25^, apparent *J*_*max*_^25^ and *E*_*a*_) which did not met the assumptions of residual normality and homoscedasticity even with transformations. Means were compared by the adjusted Tukey method and differences were considered significant if *P* ≤ 0.05.

## Results

### Temperature response of *A*_*n*_ and *R*_*d*_

The temperature response curve of net photosynthesis at saturated light (*A*_*n*_) followed a common parabolic shape (Fig. 1a, 1b). The two hybrid poplar clones adjusted their thermal optimum (*T*_*opt*_) of *A*_*n*_ in response to growth temperature. Low nitrogen level constrained the adjustment of *T*_*opt*_ for clone M×N but not M×B (Table 2). Also, *T*_*opt*_ was lower than growth temperature except for clone M×B at 23°C. The two hybrid poplar clones showed different trends regarding *A*_*n*_ at *T*_*opt*_ (*A*_*opt*_) which increased with increasing growth temperature for clone M×B and remained unaffected for clone M×N. *A*_*growth*_ had a similar trend as *A*_*opt*_ in response to growth temperature and N level. Both *A*_*growth*_ and *A*_*opt*_ declined at low N level for both clones (Table 2).

The two hybrid poplar clones had different strategies in term of thermal response of dark respiration (*R*_*d*_) (Fig. 1c, 1d). Irrespective of N level, basal rate of *R*_d_ (*R*_*d*_^*10*^) decreased by augmenting growth temperature for clone M×B but remained unchanged for clone M×N (Table 2). The rate of change in *R*_d_ by 10°C change in temperature, *Q*_*10*_, decreased when growth temperature was increased, irrespective of *N* level for clone M×N. In contrast, *Q*_*10*_ of clone M×B increased in response to growth temperature raise when N level was high and unchanged at low N level (Table 2).

**Fig 1.**
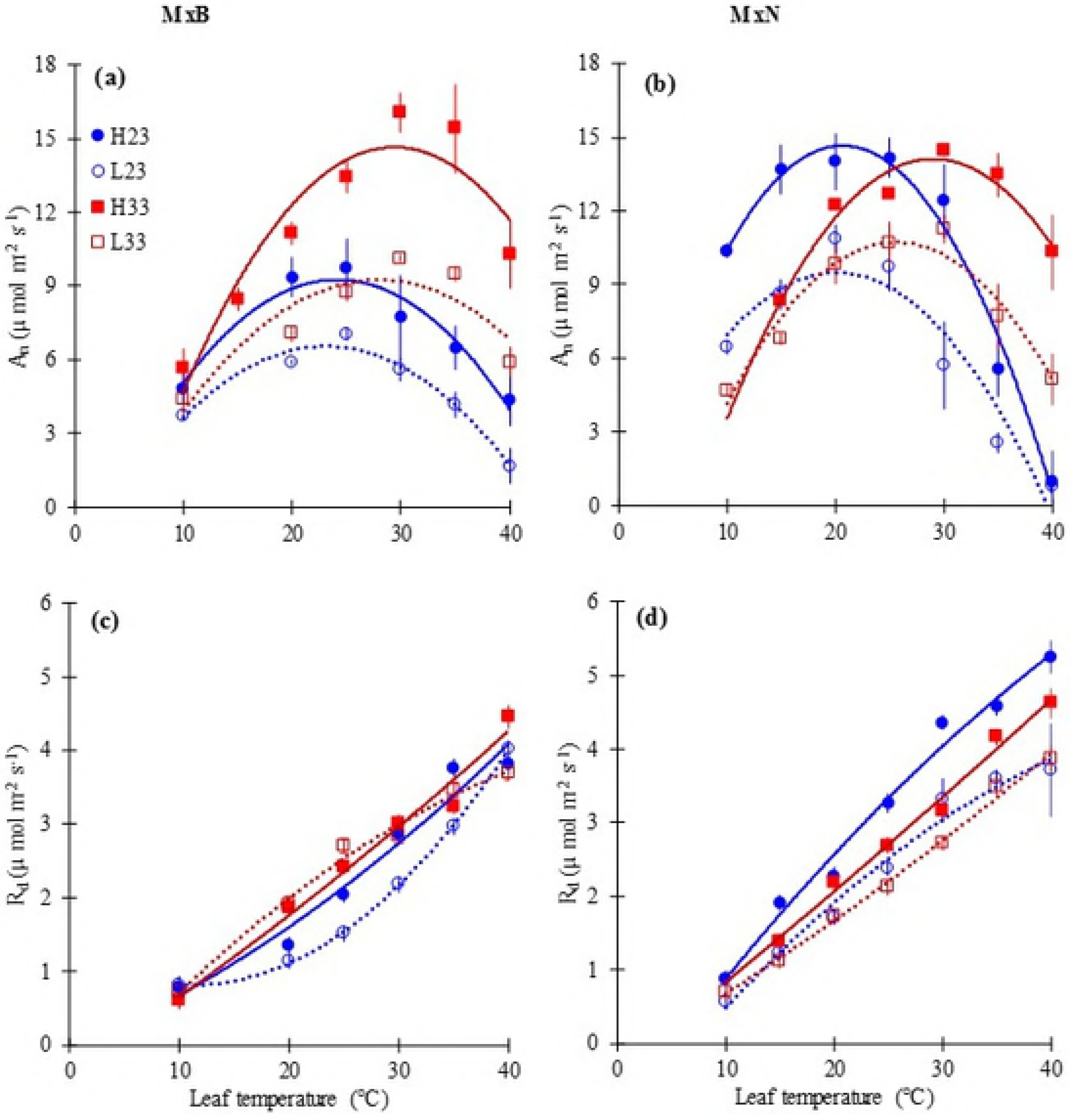
Response of net photosynthesis (*A*_*n*_) and dark respiration (*R*_*d*_) to leaf temperature for hybrid poplar clone M×B (a, c) and clone M×N (b, d) grown under two temperatures and two nitrogen levels.

H23 and L23 are treatments of high and low nitrogen level respectively at an ambient day temperature of 23°C; H33 and L33 are treatments of high and low nitrogen level at 33°C ambient day temperature. Data are represented by means ± SE (n=3). *P* value and *R*^*2*^ of curves are given in Table S1.

**Table 2:** Means (±SE) of thermal acclimation-related traits of two hybrid poplar clones (M×B and M×N) grown at day/night temperature of 23/18°C and 33/27°C under high (HN) and low (LN) nitrogen levels (n=3).

|  | Clone M×B |  |  |  | Clone M×N |  |  |  |
| --- | --- | --- | --- | --- | --- | --- | --- | --- |
|  | 23°C |  | 33°C |  | 23°C |  | 33°C |  |
|  | HN | LN | HN | LN | HN | LN | HN | LN |
| $T_{opt}(A_n)$ | 23.1 (1.2)<sup>bc</sup> | 24.1 (1.2)<sup>b</sup> | 30.3 (1.3)<sup>a</sup> | 29.3 (1.3)<sup>a</sup> | 20.5 (1.2)<sup>c</sup> | 19.7 (1.3)<sup>c</sup> | 30.1 (1.3)<sup>a</sup> | 26.1 (1.3)<sup>b</sup> |
| $A_{opt}$ | 10.1 (0.8)<sup>b</sup> | 7.1(0.8)<sup>d</sup> | 14.9 (0.8)<sup>a</sup> | 9.3 (0.8)<sup>cb</sup> | 15.1 (0.8)<sup>a</sup> | 9.1 (0.8)<sup>c</sup> | 14.2 (0.8)<sup>a</sup> | 10.9 (0.8)<sup>b</sup> |
| $A_{growth}$ | 10.6(1.0)<sup>b</sup> | 6.9(0.9)<sup>c</sup> | 13.9(1.1)<sup>a</sup> | 8.9(1.1)<sup>c</sup> | 14.9(1.1)<sup>a</sup> | 9.6(0.9)<sup>c</sup> | 13.4(1.1)<sup>a</sup> | 9.5(1.1)<sup>c</sup> |
| $Rd_{10}$ | 0.76(0.1)<sup>b</sup> | 0.80(0.1)<sup>ab</sup> | 0.60(0.1)<sup>cd</sup> | 0.70(0.1)<sup>c</sup> | 0.90(0.1)<sup>a</sup> | 0.56(0.1)<sup>d</sup> | 0.90(0.1)<sup>a</sup> | 0.71(0.2)<sup>cd</sup> |
| $Q_{10}(Rd)$ | 1.9(0.1)<sup>b</sup> | 1.8(0.1)<sup>b</sup> | 2.0(0.1)<sup>a</sup> | 1.9(0.1)<sup>b</sup> | 2.0(0.1)<sup>a</sup> | 2.2(0.1)<sup>a</sup> | 1.8(0.1)<sup>b</sup> | 1.8(0.1)<sup>b</sup> |
| $g_{s\_growth}$ | 0.16(0.01)<sup>c</sup> | 0.17(0.01)<sup>bc</sup> | 0.26(0.01)<sup>a</sup> | 0.20(0.01)<sup>b</sup> | 0.20(0.01)<sup>b</sup> | 0.16(0.01)<sup>c</sup> | 0.18(0.01)<sup>cb</sup> | 0.13(0.01)<sup>d</sup> |
| $V_{cmax}^{25}$ | 53(6)<sup>c</sup> | 34(6)<sup>d</sup> | 70(6)<sup>ab</sup> | 38(6)<sup>d</sup> | 78(6)<sup>a</sup> | 54(6)<sup>bc</sup> | 62(7)<sup>b</sup> | 53(6)<sup>bc</sup> |
| $J_{max}^{25}$ | 146(7)<sup>b</sup> | 67(7)<sup>d</sup> | 106(6)<sup>c</sup> | 74(9)<sup>d</sup> | 240(9)<sup>a</sup> | 124(7)<sup>c</sup> | 162(8)<sup>b</sup> | 107(8)<sup>c</sup> |
| $T_{opt}(V_{cmax})$ | 33(1.5) | - | NA | NA | 34(1.2) | - | NA | NA |
| $E_a(V_{cmax})$ | 75 (3)<sup>a</sup> | - | 49(3)<sup>b</sup> | 57(3)<sup>b</sup> | 58(3)<sup>b</sup> | - | 54(3)<sup>b</sup> | 55(3)<sup>b</sup> |
| $T_{opt}(J)$ | 34 (1.9) | - | NA | NA | 30(1.1) | - | NA | NA |
| $E_a(J)$ | 46(2)<sup>a</sup> | - | 32(2)<sup>b</sup> | 28(2)<sup>b</sup> | 34(2)<sup>ab</sup> | - | 28(2)<sup>b</sup> | 33(2)<sup>b</sup> |
| $J_{max}^{25}:V_{cmax}^{25}$ | 2.43(0.1)<sup>ab</sup> | 1.68(0.15)<sup>c</sup> | 1.53(0.15)<sup>c</sup> | 1.81(0.19)<sup>bc</sup> | 2.68(0.15)<sup>a</sup> | 2.01 (0.15)<sup>bc</sup> | 2.42(0.19)<sup>ab</sup> | 1.95(0.16)<sup>b</sup> |
| $SLA$ | 172(7)<sup>a</sup> | 132(7)<sup>cd</sup> | 154(8)<sup>b</sup> | 123(7)<sup>d</sup> | 143(7)<sup>b</sup> | 141(7)<sup>bc</sup> | 136(8)<sup>c</sup> | 112(8)<sup>e</sup> |
| $N_{area}$ | 1.3(0.1)<sup>b</sup> | 0.8(0.1)<sup>d</sup> | 1.3(0.1)<sup>b</sup> | 0.8(0.1)<sup>d</sup> | 1.9(0.1)<sup>a</sup> | 1.1(0.1)<sup>c</sup> | 1.4(0.1)<sup>b</sup> | 1.1(0.1)<sup>c</sup> |

Within rows, means followed by the same letter do not differ significantly at $\alpha=0.05$ based on Tukey's test

### Temperature response of apparent*V*_cmax_ and *J*

Apparent *V*_cmax_^25^ was insensitive to growth temperature at low N level for both clones. In contrast, at high N level, apparent *V*_cmax_^25^ increased for clone M×B and decreased for clone M×N when growth temperature was increased (Table 2). Apparent *J*_max_^25^ decreased with increasing growth temperature for plants growing at high N level and was insensitive to growth temperature at low N level. The ratio *J*_max_^25^: *V*_cmax_^25^ decreased with increasing growth temperature at high N level for clone M×B but not for clone M×N (Table 2). At low N level, *J*_max_^25^:*V*_cmax_^25^ ratio was insensitive to growth temperature.

The temperature response curve of apparent *V*_cmax_ and apparent *J* were affected by growth temperature but not by nitrogen level. In fact, at cooler growth temperature, apparent *V*_cmax_ peaked at 33°C and 34°C (Fig. 2; Table 2) and apparent *J* peaked at 34°C and 30°C (Fig. 2; Table 2) for clones M×B and M×N respectively. However, apparent *V*_cmax_ and apparent *J* did not show any deactivation at warm temperature (Fig. 2). The activation energy (*E*_*a*_) of apparent *V*_cmax_ and *J*, decreased with increasing growth temperature for clone M×B and remained constant for clone M×N (Table 2).

**Fig 2.**
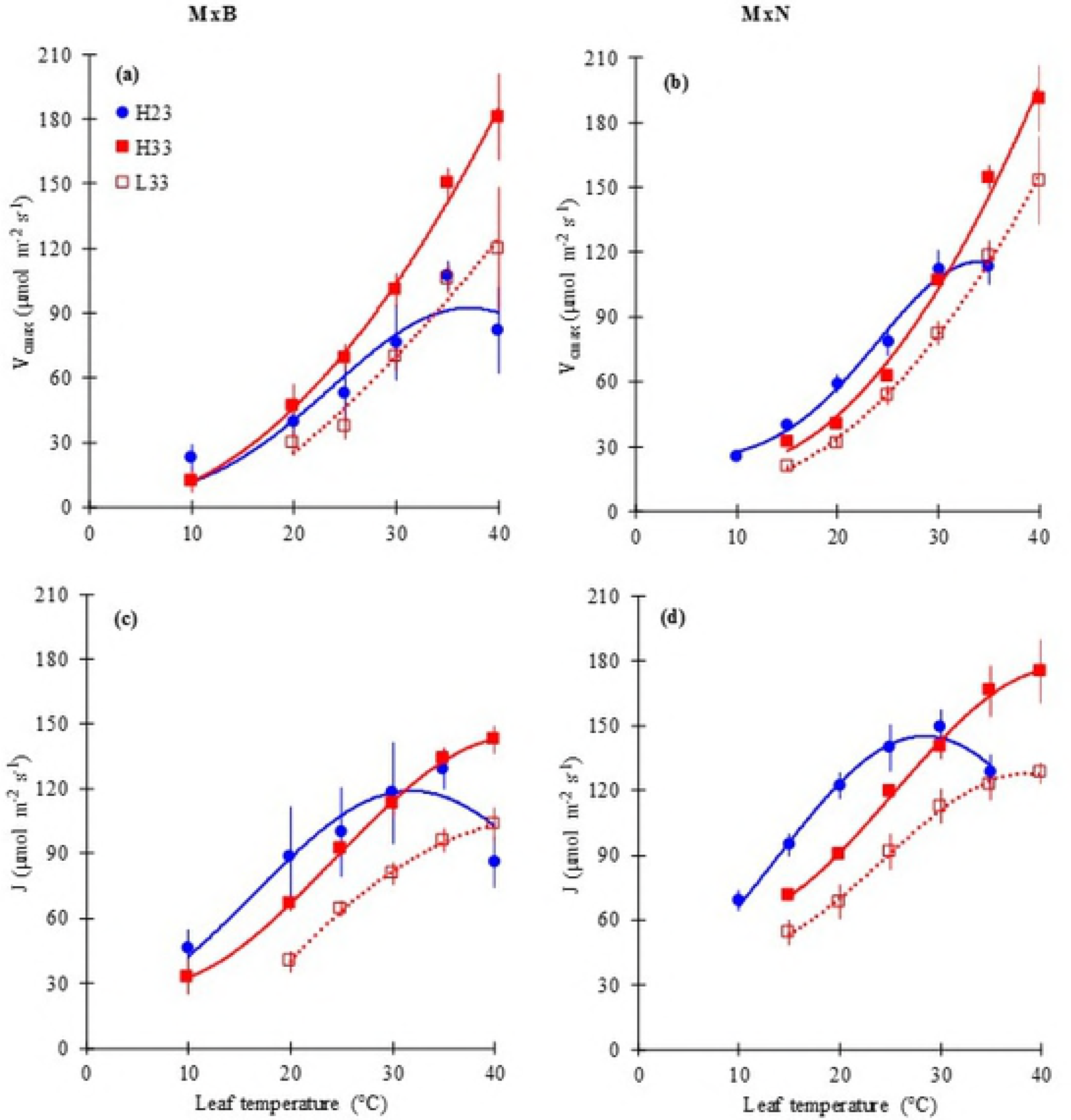
The temperature dependence of the apparent maximum carboxylation capacity of RuBisCO (*V*_cmax_) and the apparent electron transport rate (*J*) for clone M×B (a, c) and clone M×N (b, d) grown under two temperatures and two nitrogen levels.

See Fig 1 for symbols. L23 treatment was not given for both clones because A-Ci curves at 35 and 40 °C failed to converge and estimates of *V*_*cmax*_ and *J* could not be obtained. Data are represented by means ± SE (n=3). *P* value and *R*^*2*^ of curves are given in Table S1

### Temperature response of stomatal conductance(*g*_s_)

*g*_s_ decreased under all treatments and for both clones when *T*_*leaf*_ was increased over the 10-40°C gradient (Fig. 3). *g*_s_ at the growth temperature, derived from the *g*_*s*_-*T* response curves (*g*_*s_growth*_) was influenced by both clone and growth temperature. For clone M×B, *g*_*s_growth*_ was 62.5 % and 17 % higher at warm, compared to cooler growth temperature under high and low nitrogen level respectively (Table 2). Conversely, for clone M×N, *g*_*s_growth*_ was similar among growth temperature at high N level averaging 0.19 mol H_2_O m^−2^ s^−1^ and decreased by increasing growth temperature at low N level (0.16 vs. 0.13 mol H_2_O m^−2^ s^−1^).

**Fig 3.**
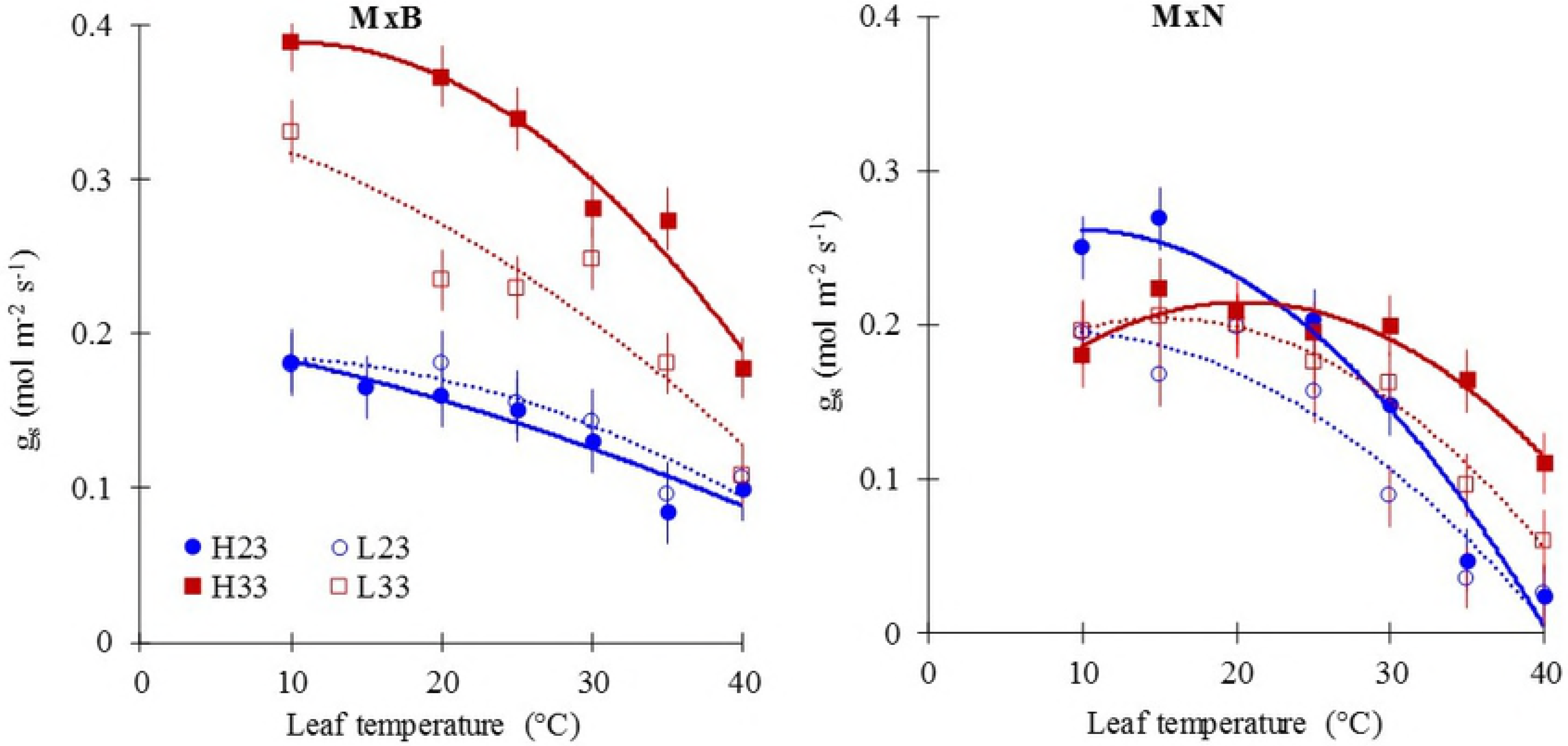
Response of stomatal conductance (*g*_*s*_) to leaf temperature of two hybrid poplar clones (M×B) and (M×N) grown under two temperatures and two nitrogen levels (n=3).

See Figure 1 for symbols. Data are represented by means ± SE (n=3). *P* value and *R*^*2*^ of curves are given in Table S1

### RuBisCO and RuBisCO activase amount

Relative amount of RuBisCO (*RAR*) decreased significantly when N level changed from high to low (Fig. 4a). *RAR* did not change in response to change of growth temperature for both clones (Fig. 4a). In addition, at high N level, *RAR* was similar between clones, being around 0.8 on average. At low N level, RAR was two folds higher for clone M×N compared to clone M×B (Fig. 4a). Nitrogen enrichment remarkably increased the relative amount of RuBisCO activase (*RARCA*), particularly for clone M×N which had a lower *RARCA* at low N level, compared to M×B (Fig. 4b). Except for clone M×B at high N, *RARCA* was stimulated by warmer growth temperature (Fig. 4b). More importantly, the ratio of short isoform to large isoform of *RCA* was markedly simulated by warm conditions for clone M×N and only at low N for clone M×B (Fig. 5).

**Figure 4:**
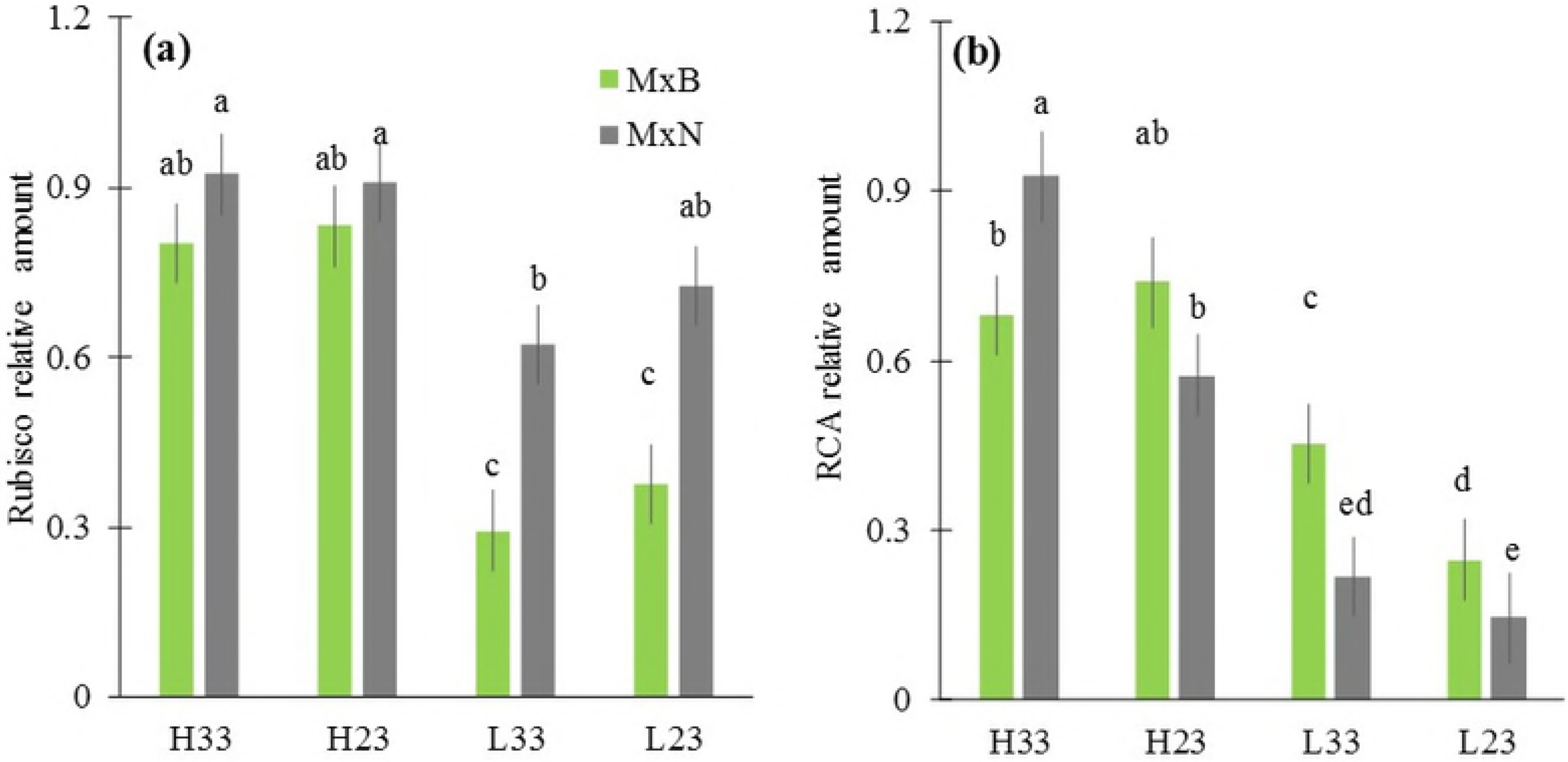
Relative amounts of RuBisCO (a) and RuBisCO-activase (b) measured by western blot for two hybrid poplar clones (M×B) and (M×N) grown under two temperatures and two nitrogen levels (n=3).

Proteins were extracted from leaves and analysed by SDS-PAGE. Immunoblots were probed with anti-Rubisco or anti-RCA antibody. H23 and L23 are treatments of high and low nitrogen level respectively at 23°C ambient daytime temperature; H33 and L33 are treatments of low and high nitrogen level at 33°C ambient daytime temperature. Data are represented by means ± SD (n=3). Means having the same letters are not significantly different at α= 0.05 based on Tukey’s tests.

**Figure 5:**
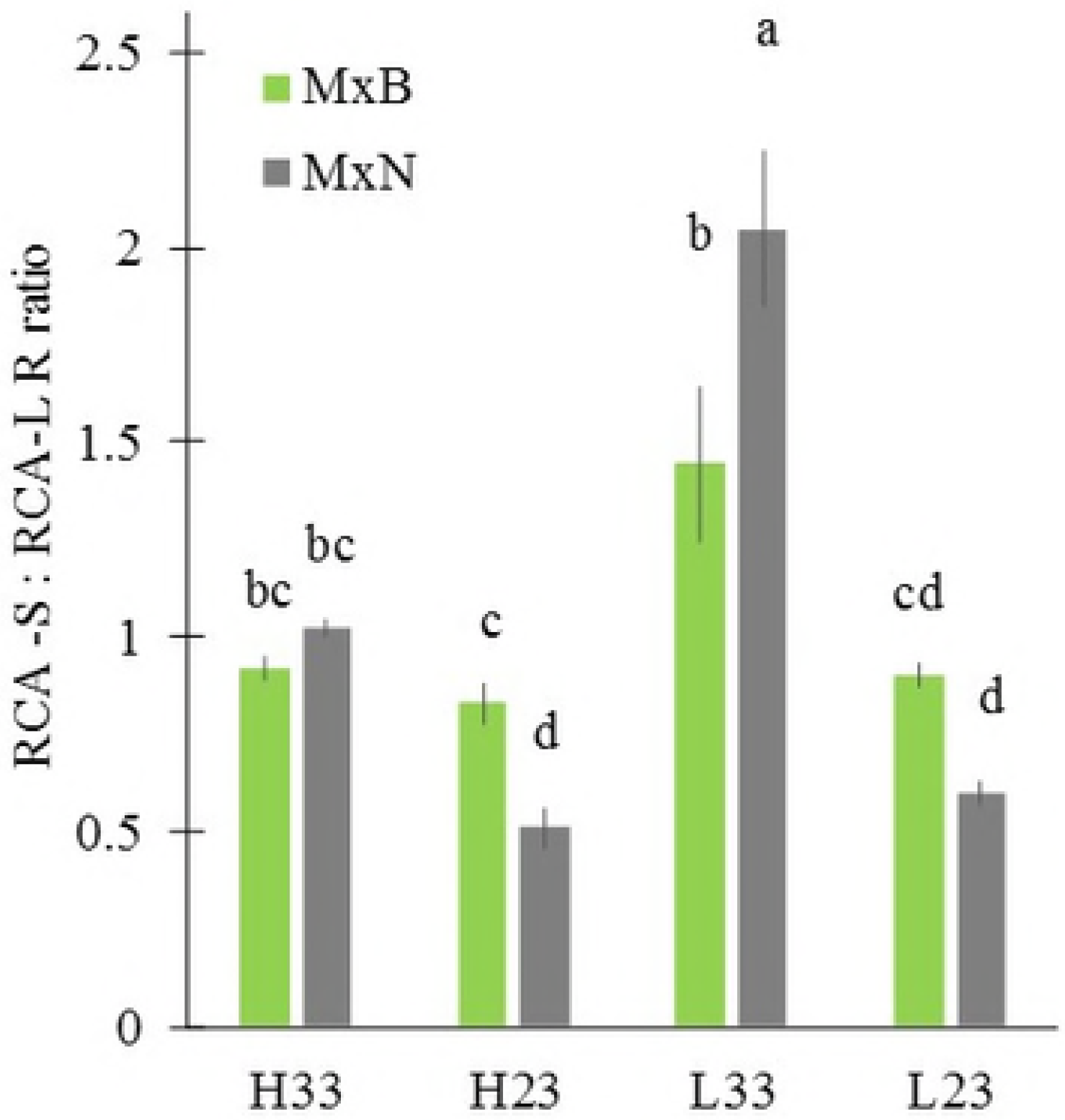
Ratio of short to long isoform of RuBisCO-activase (RCA) of two hybrid poplar grown under two temperatures and two nitrogen levels (n=3).

RCA-S: short isoform of RCA; RCA-L: long isoform of RCA. Data are represented by means ± SE (n=3). Means having the same letters are not significantly different at α= 0.05 based on Tukey’s tests.

## DISCUSSION

### Thermal acclimation of *A*_*n*_ and *R*_d_

The two hybrid poplar clones showed a clear thermal acclimation of *A*_*n*_ by adjusting *A*_*opt*_ and/or *T*_*opt*_ to growth tempearture. This is in accordance with results of Silim et al. (2010) on cold and warm ecotypes of *Populus balsamifera* which maintained *A*_*opt*_ without an evident change of *T*_*opt*_. We found that *T*_*opt*_ of *A*_*n*_ under warm tempearture was identical to mean growth temperature (the average of day time/night-time =30°C) and was 3°C below the daytime growth temperature (33°C) suggesting a partial acclimation of photosynthesis rate if we assume the latter was unrelated to night-time temperature. So far, studies focusing on night-time temperature effect on *A*_*n*_ are very scarce (Turnbull et al., 2002). *T*_*op*_ of *A*_*n*_ for clone M×N was lowered by low nitrogen level under warms conditions. This result is difficult to explain from traits measured in our study and could be an outcome of a differential expression of proteins. The net photosynthetic rates at the growth temperature (*A*_*growth*_), a relevant quantitative trait of thermal acclimation of *A*_*n*_ (Way and Yamori, 2014; Yamori et al., 2014), was enhanced in plants grown at the warm temperature for clone M×B and remained unchanged for clone M×N. These results suggest a differential thermal adaptation range of the two hybrid poplar clones which could result from the climate of origin of their parents. Hence, the choice of suitable clones based on their thermal acclimation capacity would increase productivity of hybrid poplar plantations under future warming conditions, particularly in heat-prone regions like the south of Québec, Canada.

Thermal acclimation of *R*_d_ is very common for C_3_ plants and several studies reported a downshift of *R*_*d*_^*10*^ (Type II acclimation) and a decrease of *Q*_*10*_ (Type I acclimation) in reponse to warmer temperatures (Atkin and Tjoelker, 2003; Reich et al., 2016) but few studies on *Populus* exist in this regard (Dillaway and Kruger, 2011; Ow et al., 2008; Silim et al., 2010; Tjoelker et al., 1999). In accordance with the finding of Tjoelker et al. (1999) for *Populus tremula*, we found substantial Type I acclimation of *R*_*d*_ (downshift of *Q*_*10*_) to growth temperature for clone M×N. In contrast, no acclimation of *R*_*d*_ was observed for clone M×B which may be related to the unchanged density of mitochondria (Atkin and Tjoelker, 2003). Moreover, nitrogen had no effect on thermal acclimation of *Q*_*10*_ for both clones as observed in other tree species (Benomar et al., 2018; Tjoelker et al., 1999).

### Thermal response of photosynthetic biochemical limitations

The effect of growth temperature on temperature response curve of apparent *V*_cmax_ and *J* in terms of their values at reference temperature of 25°C, their *T*_*opt*_ and their activation energy is species-dependant as reported by recent studies (Benomar et al., 2018; Hikosaka et al., 2006; Hikosaka et al., 1999; Kattge and Knorr, 2007; Slot and Winter 2017; Way and Yamori 2014). In our study, the apparent *V*_cmax_^25^ stimulated by warm growth temperature for clone M×B, might explain the noticeable increase of *A*_*opt*_ (up to 50 %) by warmer growth conditions under high N level. In parallel, the small decrease in the apparent *V*_cmax_^25^ at warm growth conditions observed for clone M×N might explain the observed similar *A*_*opt*_ under the two growth temperature. These results are in agreement with the findings of other studies showing a similar or a greater *V*_cmax_^25^ when growth temperature increased (Aspinwall et al., 2017; Silim et al., 2010; Way and Yamori, 2014; Slot and Winter, 2017). In contrast, the apparent *J*_max_^25^ decreased at warmer growth temperature as reported for *Populus balsamifera* (Silim et al., 2010) and other tree species (Yamori et al., 2008; Way and Yamori, 2014; Slot and Winter, 2017). Hikosaka et al. (2006) suggested the increase in the activation energy of *V*_*cmax*_ (*Ea*) with the increase in growth temperature as an explanatory mechanism of thermal acclimation of *A*_*n*_ (at least by the increase of *T*_*opt*_ with growth temperature). Our results are diverging with this postulate since we obseved no change in *E*_*a*_ for clone M×N and a remarkable decrease of *E*_*a*_ for clone M×B. However, the patterns we observed have been reported for several species including *Populus tremuloides* (Dillaway and Kruger, 2010), *Populus balsamifera* (Silim et al., 2010) and *Corymbia calophylla* (Aspinwall et al., 2017).

The temperature optimum (*T*_*opt*_) of apparent *V*_cmax_ and *J* acclimated to growth temperature (Fig. 2) as observed for others species (Kattge and Knorr, 2007; Crous et al., 2018) and may have contributed in the observed acclimation of *A*_n_ (Fig. 1). Under cooler conditions, *T*_*opt*_ of apparent *V*_cmax_ and *J* were similar but much higher than that of *A*_n_ indicating a very likely involvement of other traits in the observed value of *T*_*opt*_ of *A*_*n*_ under this condition (23 °C).

The adjustment of leaf nitrogen invested in soluble *vs*. insoluble proteins in response to change in growth temperature, inferred from *J*_max_^25^ to *V*_cmax_^25^ ratio, can be achieved through the maintenance of an optimal balance between the rate of photosynthetic carboxylation *vs*. RuBP regeneration. This mechanism allows plants to maximize the photosynthetic rate at a given growth temperature (Hikosaka et al., 1999; Kattge and Knorr, 2007). Therefore, the decrease of *J*_max_^25^:*V*_cmax_^25^ ratio consequent to an increase of growth temperature has been reported to significantly contribute to thermal acclimation of *A*_*n*_ (Kattge and Knorr, 2007; Slot and Winter 2017; Crous et al., 2018). In our study, this pattern occurred for clone M×B under high N level which has resulted in an increase of both *V*_cmax_^25^ and *A*. Conversely, the lack of modulation of *J*_max_^25^:*V*_cmax_^25^ ratio for clone M×N may have contributed to the observed decrease in *V*_cmax_^25^ and to the maintenance of *A*_*opt*_. Under low N level, *A*_*opt*_ of M×B increased under warmer conditions without any change of the *J*_max_^25^:*V*_cmax_^25^ ratio. Therefore, the increase of *V*_cmax_^25^ and *A*_*opt*_ under the warm growth temperature cannot be attributed only to the shift in *J*_max_^25^:*V*_cmax_^25^ ratio.

### RuBisCO and RuBisCO activase amounts in response to experimental warming

The RuBisCO content in our study was quite sensitive to nitrogen level but not to growth temperature. Neither thermal acclimation of *A*_*n*_ (*T*_*opt*_ and *A*_*opt*_) nor *J*_max_^25^:*V*_cmax_^25^ ratio was affected by RuBisCO content. The absence of any effect of RubisCO content on traits related to thermal acclimation of *A*_*n*_ has been reported by Weston et al. (2007) and Kruse et al. (2017), while other studies found a significant decrease of *V*_*cmax*_^*25*^ linked to a decrease in RuBisCO and leaf nitrogen content (Scafaro et al., 2016; Crous et al., 2018). Thus, the relationship between the change in RuBisCO content in response to growth temperature and thermal acclimation of *A*_*n*_ via the modulation of photosynthetic capacity attributes (*V*_*cmax*_^*25*^ and *J*_*max*_^*25*^) is, most likely, depending on species and environemental parameters (e.g. nitrogen availibility). Indeed, CO_2_ conductance, the variation of *RCA* content and the temperature dependency of Rubisco kinetic properties have been rerported to be determinant factors of the *V*_cmax_^25^ response to growth temperature and consequently thermal acclimation of *A*_*n*_ (Perdomoetal et al., 2017; Way and Yamori, 2014;Yamori et al., 2006; Qiu et al., 2017). The increase of leaf *RCA* amount by increased growth temperature has been reported for several tree species (Crafts-Brandner and Salvucci, 2000; Hozain et al., 2009; Law et al., 2001; Ristic et al., 2009; Weston et al., 2007; Yamori and von Caemmerer, 2009). In our study, the hypothesized increase of *RCA* at warmer growth temperature was observed, except for clone M×B at high N. More importantly, our results demonstrated that the increase of the amount of *RCA* under warm conditions resulted mainly from increased synthesis of the short isoform which indicates that the two isoforms operate at different temperature optima.

### Stomatal conductance

The contribution of diffusional limitations to thermal acclimation of *A*_n_ remain non-well quantified for several species, including *Populus*. Our results demonstrate that the modulation of *g*_s_ (the shape of the relationship between *g*_s_ and *T*_*leaf*_ and the value of *g*_*s*_ at growth temperature) in response to change in growth temperature contributed to the observed thermal acclimation of *A*_n_ (Figure 3) as observed by Aspinwall et al. (2017), Silim et al. (2010) and Slot and Winter (2017). Also, our results suggest that the stomatal acclimation to growth temperature may be clone-specific and may have a significant impact on clone response to warming depending on soil water status. The CO_2_ diffusion in the mesophyll shares the same pathways of water transport from mesophyll to the atmosphere (Ethier et al., 2004; Flexas et al., 2013) and may lead to similar response of stomatal and mesophyll conductance to growth conditions. Moreover, a link between mesophyll conductance (*g*_*m*_) and hydraulic conductance has been reported as well (Flexas et al., 2013; Théroux-Rancourt et al., 2014), suggesting that the observed response of *g*_s_ to growth temperature may have originated from a modulation of *g*_m_ and hydraulic functioning.

In conclusion, the observed thermal acclimation of photosynthesis under our experimental conditions was clearly related to the modulation of photosynthetic capacity and *g*_*s*_ in response to growth temperature. The modulation of the photosynthetic capacity was mainly linked to *RCA* but not RuBisCO content. Further investigation regarding the involvement of mesophyll conductance and hydraulic conductivity should clarify the mechanistic basis of the observed trends.

## Acknowledgements

We thank Dr. G Ethier for his valuable comments on A-Ci curve analysis. We also thank F Larochelle and M Coyea (Université Laval) for their technical assistance throughout the project.

